# Optimizing insect metabarcoding using replicated mock communities

**DOI:** 10.1101/2022.06.20.496906

**Authors:** Elzbieta Iwaszkiewicz-Eggebrecht, Emma Granqvist, Mateusz Buczek, Monika Prus, Tomas Roslin, Ayco J.M. Tack, Anders F. Andersson, Andreia Miraldo, Fredrik Ronquist, Piotr Łukasik

**Author notes:** equal senior authorship (Fredrik Ronquist, Piotr Łukasik). Correspondence author: Elzbieta Iwaszkiewicz-Eggebrecht, Box 50007, SE-104 05 Stockholm, Sweden.

## Abstract

Metabarcoding (high-throughput sequencing of marker gene amplicons) has emerged as a promising and cost-effective method for characterizing insect community samples. Yet, the methodology varies greatly among studies and its performance has not been systematically evaluated to date. In particular, it is unclear how accurately metabarcoding can resolve species communities in terms of presence-absence, abundances, and biomass. Here we use mock community experiments and a simple probabilistic model to evaluate the performance of different metabarcoding protocols. Specifically, we ask four questions: (Q1) How consistent are the recovered community profiles across replicate mock communities?; (Q2) How does the choice of lysis buffer affect the recovery of the original community?; (Q3) How are community estimates affected by differing lysis times and homogenization?; and (Q4) Is it possible to obtain adequate species abundance estimates through the use of biological spike-ins? We show that estimates are quite variable across community replicates. In general, a mild lysis protocol is better at reconstructing species lists and approximate counts, while homogenization is better at retrieving biomass composition. Tiny insects are more likely to be detected in lysates, while some tough species require homogenization to be detected. Results are less consistent across biological replicates for lysates than for homogenates. Some species are associated with strong PCR amplification bias, which complicates the reconstruction of species counts. Yet, with adequate spike-in data, species abundance can be determined with roughly 40% standard error for homogenates, and with roughly 50% standard error for lysates, under ideal conditions. In the latter case, however, this often requires species-specific reference data, while spike-in data generalizes better across species for homogenates. We conclude that a non-destructive, mild lysis approach shows the highest promise for presence/absence description of the community, while also allowing future morphological or molecular work on the material. However, homogenization protocols perform better for characterizing community composition, in particular in terms of biomass.

## Introduction

Insects are likely the most species-rich group of animals on Earth (Stork, 2018), and fulfill a myriad of different ecosystem functions (Jordan et al., 2021). Yet our knowledge about insect diversity in natural ecosystems is still surprisingly scarce – and the key reason for this deficit is their actual diversity. Recent reports on insect abundance and diversity declines (Hallmann et al., 2017; Klink et al., 2020; Seibold et al., 2019) make monitoring more important than ever. Nonetheless, surveys based on traditional taxonomic work demand enormous time and labor. In one of the most ambitious projects aiming to characterize the insect fauna at the national level – the Swedish Malaise Trap Project – sorting 1919 samples into 300 taxonomic fractions took 15 years, despite the high manpower invested (estimated 98K person-hours devoted to sorting) (Karlsson et al., 2020). While taxonomic work remains crucial, there is a pressing need for the development of complementary high-throughput and cost-effective methods that enable adequate description of insect community diversity, composition, and spatio-temporal dynamics.

In this context, the ever-decreasing costs of high-throughput sequencing (HTS) have made DNA-based methods particularly attractive for large-scale insect biomonitoring. A number of such methods have been proposed for and tested on bulk invertebrate samples (Kennedy et al., 2020). The method perhaps most widely used today is community-level metabarcoding: the generation of a large number of barcodes for bulk samples. In this approach, DNA is first extracted from the whole multi-species sample, such as a Malaise trap catch. After this, a short region of a marker gene, typically mitochondrial cytochrome oxidase I (COI), is amplified by PCR with broad-spectrum primers and subsequently sequenced. The resulting collection of marker gene sequences – barcodes – is compared against reference databases, providing information about the identities of species present in the original bulk sample. This rapid and affordable workflow has been used to describe arthropod communities (Cristescu, 2014; Elbrecht et al., 2017; Taberlet et al., 2012) and is increasingly proving useful in ecological studies (Beng et al., 2016; Liu et al., 2020; Porter et al., 2019). Nevertheless, despite its obvious advantages, metabarcoding is still a relatively young method and many challenges remain. Perhaps most importantly, it is unclear how accurately metabarcoding can describe species diversity (presence/absence) and community composition (species’ relative abundances) – and how its accuracy can be improved.

Research to date has pointed to multiple obstacles to obtaining accurate estimates of both species-specific incidence – adding up to species diversity – and abundance from metabarcoding. Differences in species’ biomass and physical structure cause unequal contribution of DNA to the sequenced pool. Added to this, PCR errors and primer biases towards certain taxonomic groups can obscure estimates of species diversity and abundances, as demonstrated through metabarcoding of samples with known composition (Braukmann et al., 2019; Elbrecht and Leese, 2015; Martoni et al., 2022). Multiple solutions have been proposed to address these challenges. To account for differential effects of the DNA extraction procedure and later processing steps on the sequencing yields, several studies have proposed the introduction of known amounts of either tissue or DNA (so called spike-ins) (Ji et al., 2020; Lamb et al., 2019; Thomas et al., 2016) to facilitate normalizing sequencing results between and within samples.

Many crucial considerations in metabarcoding relate to early stages of sample processing – before the DNA extraction. At present, the most common way of treating bulk samples is to homogenize specimens into an “insect soup” (Elbrecht et al., 2017; Liu et al., 2020; Morinière et al., 2016; Yu et al., 2012). This destructive method results in high DNA yields, but also in the loss of all morphological information – a discouraging outcome for taxonomists and researchers dealing with irreplaceable samples. Homogenization can also decrease the odds of detection of smaller and/or rare insects, since large or abundant species will contribute much more DNA to the final pool and thereby swamp the DNA of smaller and/or rare species during sequencing. An alternative approach is then to use a non-destructive mild lysis protocol, in which insects soak in the lysis buffer for a short period. The procedure leaves specimens almost intact – as demonstrated in soft-bodied springtails (Collembola) by Porco and collaborators (2010) and in a diverse arthropod sample by Marquina and collaborators (2022) – allowing future taxonomic work. The mild lysis approach has been developed and applied in freshwater and terrestrial arthropod studies (Batovska et al., 2021; Carew et al., 2018; Martins et al., 2019; Nielsen et al., 2019), showing promising results in terms of evenness in species representation and detection. However, critical methodological parameters such as the lysis buffer composition, or lysis duration and conditions, have not been systematically investigated – despite their likely effects on the relative DNA yield from different species, and thus on any estimates of community composition.

Arguably, the best way of evaluating methodological choices in metabarcoding is their application to artificial communities of known composition, also called mock communities (Cristescu, 2014). Here, by controlling specimen numbers and approximate biomass, we obtain the reference that any metabarcoding results can be compared against. Since metabarcoding is sensitive to multiple sources of variation, making sufficient replication is the key to obtaining robust results.

However, many studies to date have used wild-caught insects to construct mock communities (Batovska et al., 2021; Zizka et al., 2019), which precludes proper replication since each sample is unique. Others have based their approach on DNA rather than insect samples and constructed mock communities by blending species-specific DNA extracts, or extracts from low-diversity samples into more complex blends (Braukmann et al., 2019; Nielsen et al., 2019). While such approaches might be suitable for testing PCR and sequencing steps of the workflow, they depart significantly from realistic community sample processing procedures. We argue that well-replicated, relevant studies may best be obtained by constructing mock communities from size-standardized, homogeneous populations of different species. Such samples can be prepared in multiple copies, allowing us to evaluate the robustness of our inferences made from metabarcoding.

In this study, we use a set of individual-based mock communities as a powerful tool to systematically compare the effects of methodological choices on inferences from metabarcoding data. To scrutinize our ability to accurately describe insect communities in terms of species presence/absence, abundance and biomass, we use metabarcoding of replicate community samples to generate taxonomically-assigned sequence yields, and a simple probabilistic model to analyze the results. We revisit the impact of different methodological solutions in three experiments using mock communities, and ask: (Q1) How consistent are the composition profiles across replicate mock communities?; (Q2) How does the choice of buffer affect community recovery?; (Q3) How are community estimates affected by differing lysis times and homogenization?; and (Q4) Is it possible to obtain adequate species abundance estimates through the use of biological spike-ins?

## Materials and Methods

Our study is built on three separate experiments, all using replicate mock communities constructed from different combinations of up to 25 insect species (Fig. 1, Table S1). In Experiment I, we compare different formulae for lysis buffer, and assess if they affect the inferences of species presence. For Experiment II, we select the most promising buffer and then test a wider range of lysis times applied to larger and more diverse communities. In both Experiments I and II, we also assess how mild lysis and homogenization methods compare, and how they affect our ability to accurately reconstruct the true composition of mock communities. Finally, in Experiment III, we use biological spike-ins to measure how accurately we can recover known species abundances from bulk samples subjected to mild lysis, as well as to homogenization.

**Figure 1.**
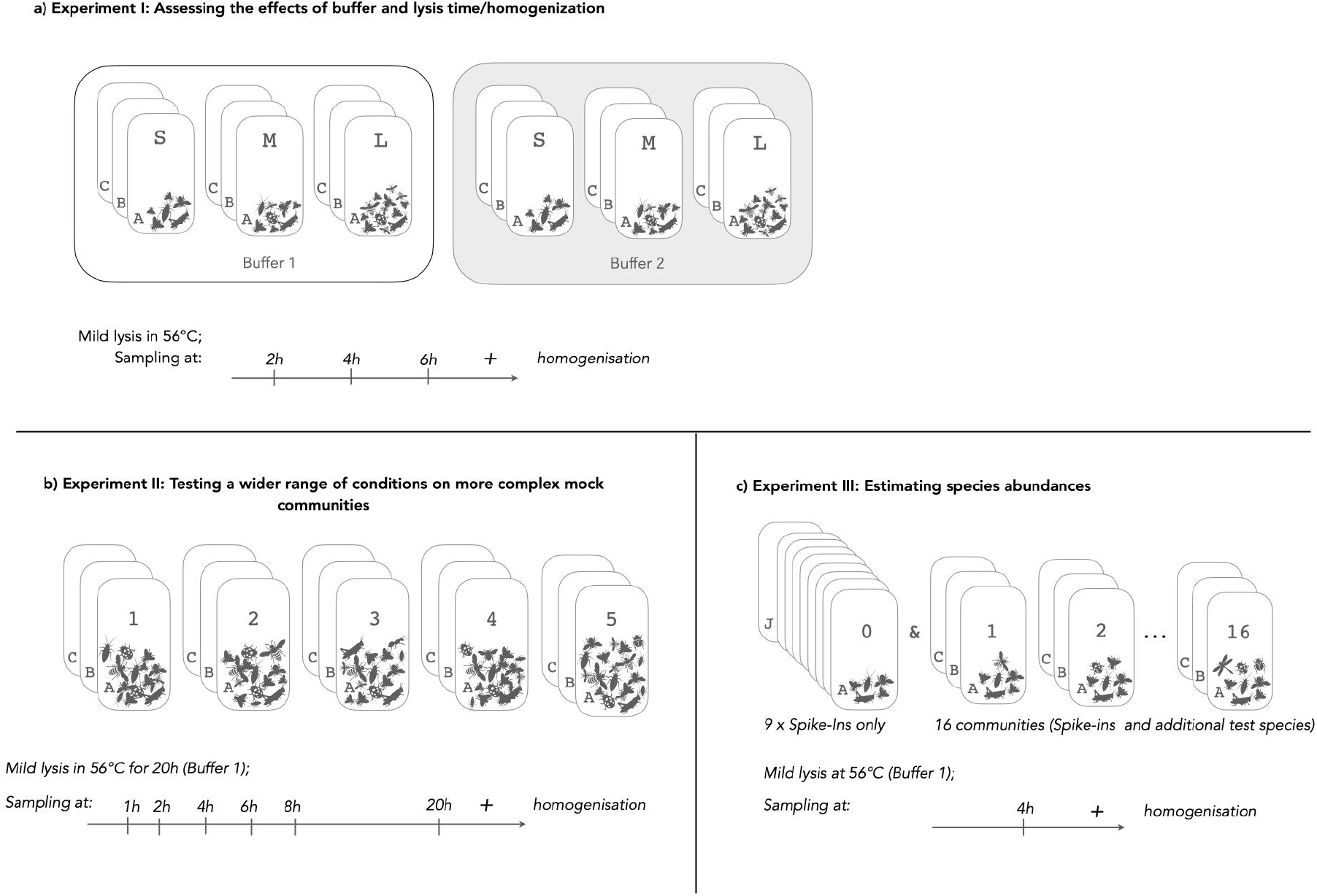
Visual summary of three Experiments that form the basis of the current study. (A) In Experiment I, we tested two lysis buffers on three mock communities of different sizes (S, M and L: up to 25 individuals of 13 species), prepared in six replicates, three of which were incubated with Buffer 1 and the other three with Buffer 2. We collected lysate aliquots at three timepoints for each replicate before homogenizing them. (B) In Experiment II, we constructed five larger and more complex communities (up to 132 individuals of 25 species), preparing each in three replicates. All samples were incubated in Buffer 1 with six lysate aliquots taken, followed by homogenization. (C) In Experiment III, we prepared nine identical communities made of five reference (spike-in) species only (“0”) and 16 different small (6-23 individuals per sample) mock communities made out of the five spike-ins and a varying number of seven other species. Communities, prepared in triplicate, were lysed for 4h and then homogenized.

### Constructing mock communities (Q1)

We constructed communities using 31 standardized, reference insect species obtained from biocontrol and pet stores, hobbyists, laboratory cultures, and social insect colonies (Table S1). All individuals within a species represented the same developmental stage and were size-standardized. Insects were preserved in 90% ethanol, and stored at −20°C until used. Species-specific dry weights were calculated as averages from ten specimens (Table S1). Communities were prepared by combining pre-set numbers of individuals from each species in a Falcon tube with ethanol. Every mock community was prepared in at least three identical copies. Each experiment used different community types, as explained below.

### Experiment I: Assessing the effects of buffer and lysis time/homogenization (Q2, Q3)

To look into the effect of different lysis buffers we assembled three mock communities - S, M and L - consisting of 8-13 species represented by 11-25 individuals (Fig. 1, Table S2). Each community was prepared in six replicates: three were lysed with Buffer 1 (Vesterinen et al., 2016; modified from Aljanabi and Martinez, 1997), and the other three with Buffer 2 (arthropod-specific buffer from the Canadian Centre for DNA Barcoding (CCDB; Ivanova et al., 2006). Communities were incubated in 5mL of the buffer, in a shaking water bath set to 56°C. 50uL aliquots were taken after 2h, 4h and 6h of incubation. Then samples were homogenized using bead beaters. DNA was purified from 20μL aliquots of the lysate or homogenate. (For buffer recipes and details see Text S1).

### Experiment II: Testing a wider range of conditions on more complex mock communities (Q3)

To further explore how mild lysis time influences community estimates, and how homogenization and mild lysis compare, we expanded the range of lysis times, and increased the size, number and complexity of mock communities. We used five distinct communities, comprising 20-25 species represented by 68-132 total individuals. Each mock community was replicated three times (Fig. 1b, Table S3). The samples were incubated in 50mL tubes with 20mL of Buffer 1 at 56°C in a shaking water bath. 250ul aliquots of lysate were taken at six time points during the incubation (1h, 2h, 4h, 6h, 8h and 20h). Afterwards, each sample was homogenized using a bead beater. A 225μL aliquot of each lysate or homogenate was then purified using silica-coated magnetic beads with the KingFisher Cell and Tissue DNA kit on a KingFisher Duo robot (both Thermo Scientific, USA).

### Experiment III: Estimating species abundance (Q4)

To explore the utility of biological spike-ins in improving abundance estimations, we assembled 17 mock communities, using up to 12 species. Five species: *Shelfordella lateralis* (Blattodea: Blattidae), *Drosophila hydei, Drosophila yakuba* (both Diptera: Drosophilidae), *Gryllus bimaculatus and Gryllodes sigillatus* (both Orthoptera: Gryllidae) were used for calibration and we will henceforth refer to them as “spike-ins”. One community type (“0”) comprised only those five spike-ins, and was prepared in nine replicates. The other sixteen communities (numbered 1-16) had a variable number of additional test species (1-18 individuals representing up to six species) and were prepared in three copies (Fig. 1c, Table S4). Communities were subjected to mild lysis in Buffer 1 at 56°C for 4h and further homogenized using bead beaters. DNA was purified as in Experiment I.

### Library preparation and bioinformatics

Amplicon libraries were prepared using a two-step PCR approach, as described in Marquina et al. (2021). We amplified a 458-bp region of the COI gene using BF3-BR2 primers (Elbrecht et al., 2019) and sequenced the library pools on Illumina MiSeq v3 (2×300bp reads). For details on molecular work, see Text S2.

We used *cutadapt v.3.2* (Martin, 2011) for primer trimming and proceeded further with amplicon analysis with the *DADA2 R* package (Callahan et al., 2016) in *R* environment v.4.0 (R Core Team, 2020). Sequences were dereplicated (*derepFastq()*), denoised (*learnErrors(), dada()*) and merged (*mergePairs()*) and chimeras were removed (*removeBimeraDenovo()*), resulting in ASV (amplicon sequence variants) count tables. Since the makeup of communities was known, ASVs other than those representing verified, error-free species sequences were removed from the count tables. Detailed code, sequences and resulting count tables are available on GitHub: https://github.com/ronquistlab/optimizing_metabarcoding_mock_communities.git.

### Data visualization and statistical analysis

For Experiments I and II, we performed nonmetric multidimensional scaling (nMDS) with the *metaMDS()* function of the *vegan* package (Oksanen et al., 2020), based on Bray-Curtis dissimilarity distance. Relative read counts from Experiment I were plotted as a heatmap with the use of *pheatmap* package (Raivo, 2019) and from Experiment II as stacked bar plots with the *ggplot2 R* package (Wickham, 2016); Code and Bray-Curtis distance matrices are available on GitHub: https://github.com/ronquistlab/optimizing_metabarcoding_mock_communities.git).

For Experiment III, we developed a hierarchical Bayesian model that allowed us to estimate the variation across specimens and species in DNA yield, and the variation across samples in PCR amplification factors from spike-in data. The model parameters learned were then used to estimate the abundance and biomass of the remaining species.

As a basis for the model, let *d_tmij_* be the *DNA yield*, the amount of template DNA extracted from an insect specimen *i* of species *t* in sample *j* of community *m*, and effectively available for PCR amplification. In the simplest case, we model *d_tmij_* using a universal gamma distribution

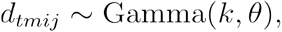

where *k* is the shape parameter and *θ* the scale parameter.

We now introduce a *PCR factor, C_j_*, specific to each sample *j*. This factor represents how many times the template DNA is multiplied in sample *j*, taking into account amplification, subsampling, sequencing depth and bioinformatic processing steps after DNA extraction. The PCR factor only affects the scale parameter of the gamma distribution, while the number of specimens or biomass affects only the shape parameter. Specifically, given that there are *n_tm_* specimens, the read count *r_tmj_* (interpreted as a concentration) is distributed as

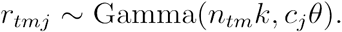

Given suitable prior probability distributions on *k*, *θ* and *c*, these parameters can be inferred from observed read counts. Specifically, we used a gamma prior on *k*, and lognormal priors on *θ* and *c*, with conjugate normal-inverse-gamma hyperpriors. For abundance estimates, we used a uniform prior on *n*. We explored variation across species in DNA yield and PCR amplification by introducing species-specific *k* and *θ* parameters, respectively.

Models were implemented as probabilistic programs in *Birch* version 1.634 (Murray and Schön, 2018), and inference was performed using sequential Monte Carlo.

Further model details and Birch code is available on GitHub: https://github.com/ronquistlab/optimizing_metabarcoding_mock_communities.git

## Results

After primer trimming and quality filtering, the sequencing yielded an average of 7573 (SD 3869) COI reads per sample (Table S5). We recovered the barcode sequences of all species used to produce the respective mock communities. However, not all species were detected in all samples where they were present. Instead the presence/absence and relative abundance of species varied substantially among mock community types, buffers (for Experiment I) and treatments (lysis time or homogenization). Furthermore, we observed major differences among replicates of the same community. We explore these patterns by answering the methodological questions defined in the Introduction.

### [Q1] How consistent are the composition profiles across replicate mock communities?

In all experiments, we observed considerable variation in the relative abundances of species, but also in their inferred presence/absence. Certain species were consistently undetected in some of the treatments (patterns explored below, Q3). For others, the rate of false negatives seemed to vary haphazardly from replicate to replicate, especially in the communities subjected to mild lysis (Figs 2, 4). As a result, the estimates of community composition varied from replicate to replicate, despite them coming from the identical mock community. In most cases, samples of one replicate lysed for different times were more similar to each other than samples lysed for the same time but originating from different replicates. The differences between replicates were becoming smaller after homogenization. For each particular lysis time, the Bray-Curtis distances between replicates were considerably higher among samples treated by mild lysis (0.22 on average in Experiment II) than between homogenized ones (0.12).

**Figure 2.**
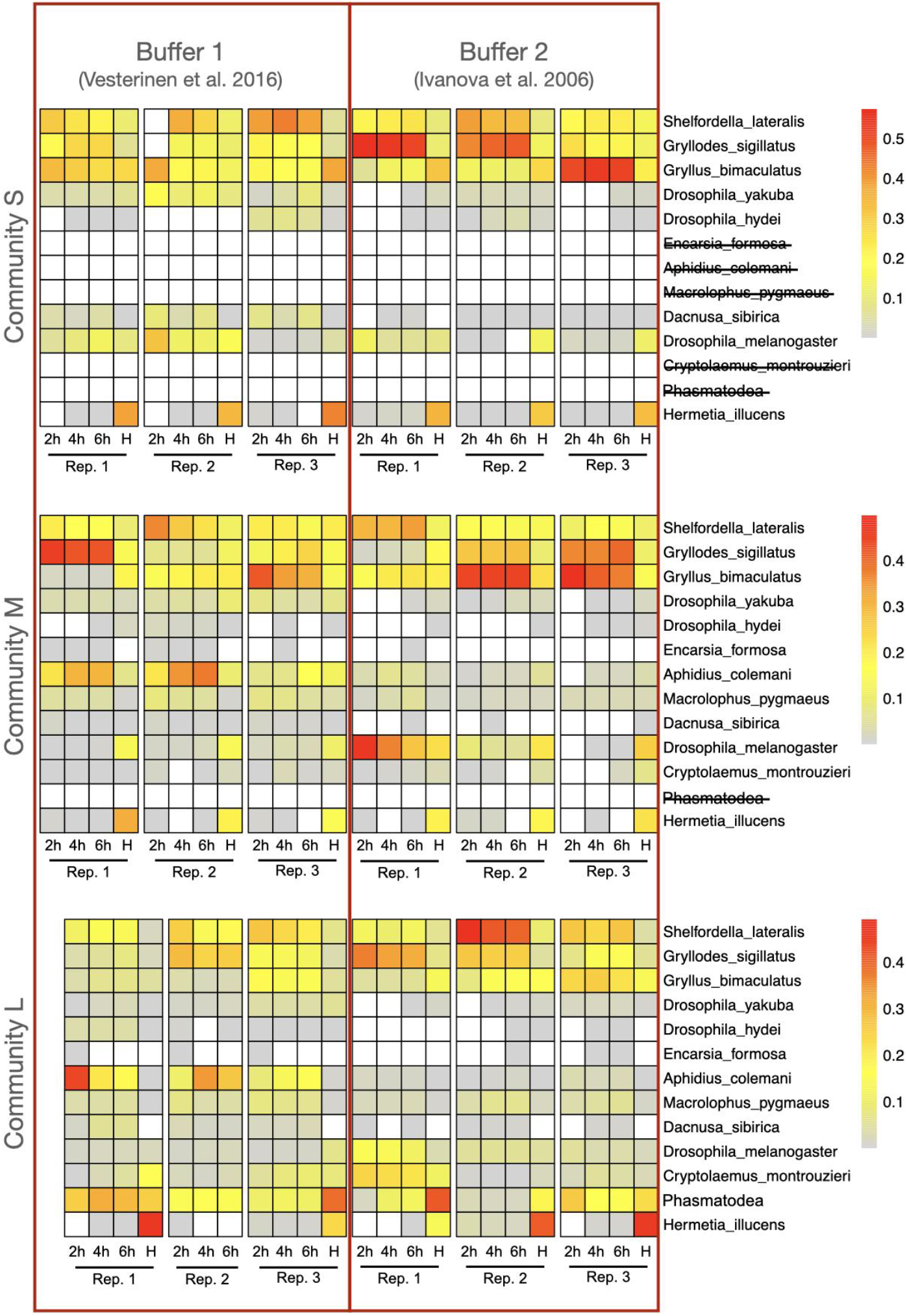
Heatmap representing relative abundance of species in Experiment I. Mock communities (three types: S, M, L) were subjected to mild lysis in two buffers (B1 and B2), with aliquots taken after 2, 4 and 6 hours, then homogenized (H). Lysate and homogenate sample aliquots were purified and used as templates for COI amplicon library preparation and sequencing. We used three biological replicates (identical copies of the mock community) per community, per buffer. Heatmap colors present relative read counts per experimental species recovered in metabarcoding. White tiles indicate 0 reads. Species not present in a given mock community have their name crossed out in the legend; note that there were no false positives.

**Figure 3.**
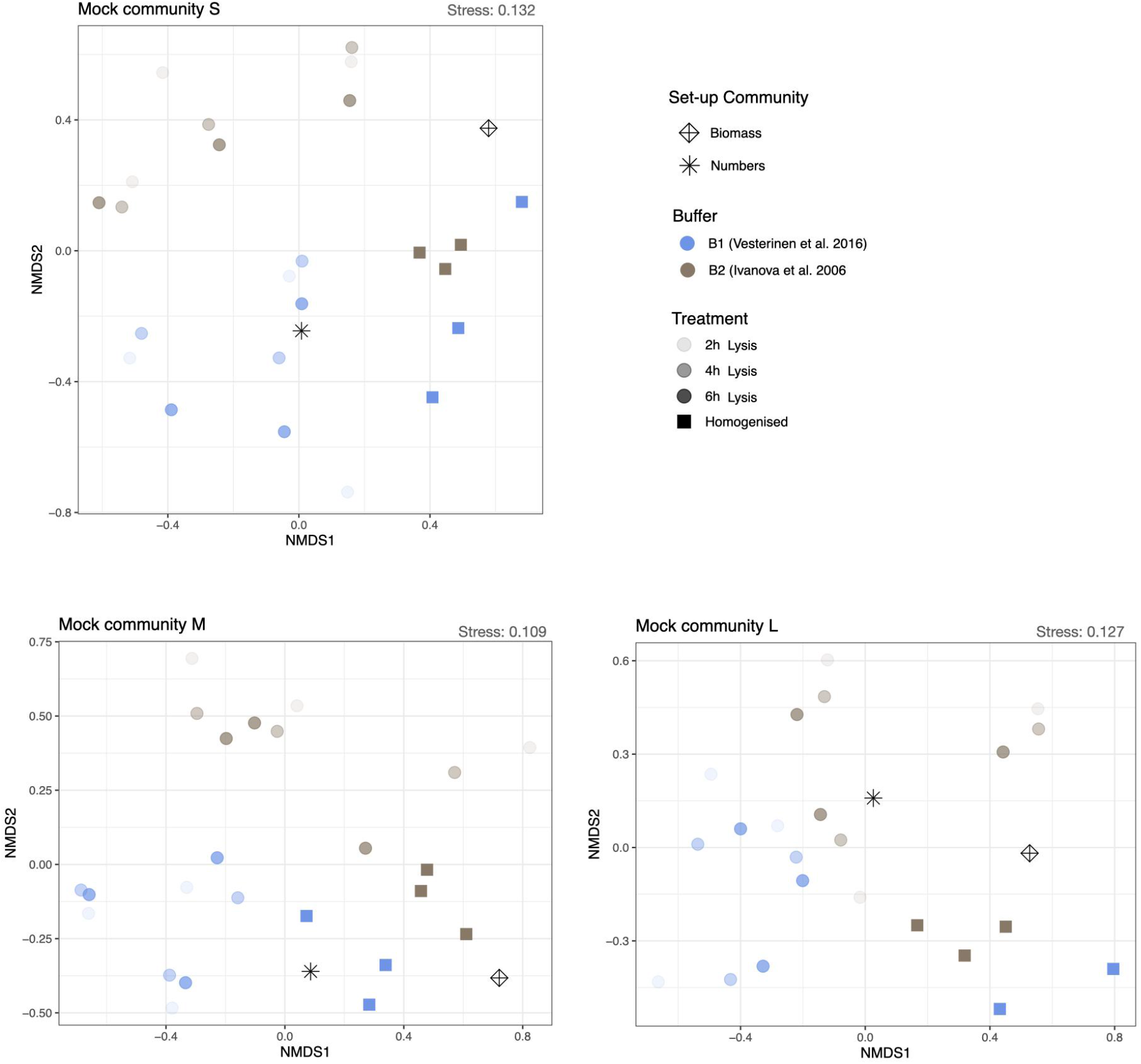
Visualization of non-metric multi-dimensional scaling (nMDS) based on metabarcoding results from Experiment I. Mock communities S, M and L are the same as in Fig. 2. Community replicates are represented as dots. Different colors represent the two buffers. Homogenized samples are squares while mild lysis are circles with lysis times represented as opacity levels. The original community set-up in terms of the relative number of species and the relative biomass of the species are represented by an asterisk and crossed diamond, respectively. Stress factor for each ordination is given in the top right corner.

**Figure 4.**
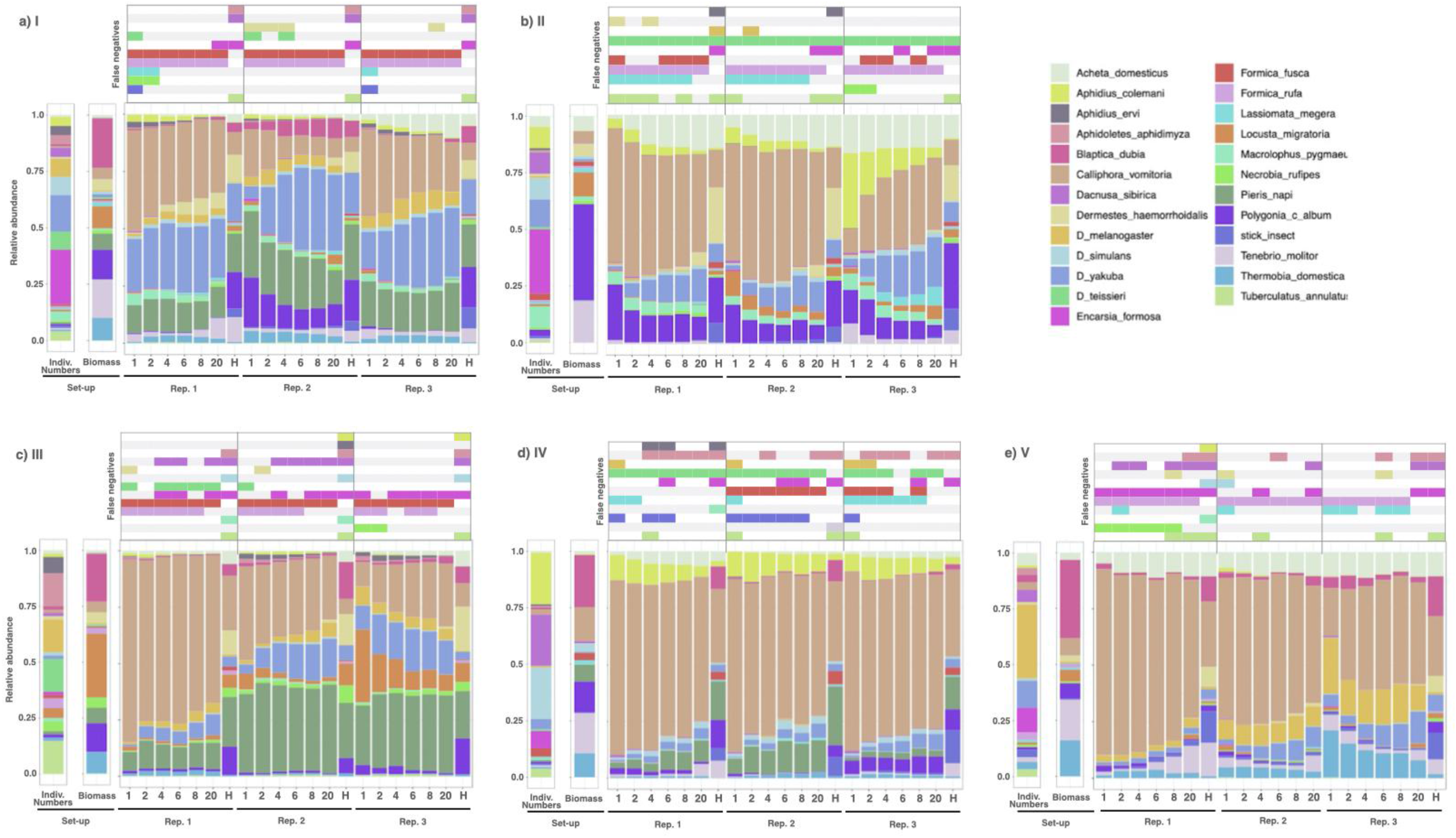
Relative species abundance in mock communities from Experiment II, based on the metabarcoding results. Panels a-e present five mock communities (I-V), each one in biological triplicate (1-3). Each panel consists of three sections: 1) two barplots showing the set-up composition of the given mock community in terms of numbers of individuals of each species (first barplot) and their relative biomass contribution to the community (second barplot); 2) figure presenting relative read counts obtained through metabarcoding for six mild lysis times and homogenates for three biological replicates one after another; 3) panel above the barplots showing false negatives, where colored fields show species that were not detected in metabarcoding even though they were present in the mock community.

### [Q2] How does the choice of lysis buffer affect community reconstruction?

In Experiment I, we found a strong effect of buffer type (Fig. 2). Overall, digestion with Buffer 1 resulted in substantially more reads from smaller species such as *Aphidius colemani, Dacnusa sibirica, Encarsia formosa*, and *Drosophila yakuba*. With Buffer 2, these small species remained undetected in multiple replicates, whereas we scored higher relative read counts from larger species such as *S. lateralis, G. sigillatus* and *G. bimaculatus* (Fig. 2).

The impact of buffer choice was further corroborated by the results of multidimensional scaling. Replicates treated by mild lysis were well separated from replicates treated by homogenization along the first axis (Fig. 3). Samples that underwent mild lysis in Buffer 1 and Buffer 2, respectively, were clearly separated from each other along the second nMDS axis, while homogenized samples clustered close to one another, regardless of the buffer used (with two individual replicates forming exceptions; see Fig. 3).

Given the improved detection of small and rare species conveyed by Buffer 1, and the more comprehensive taxonomic coverage achieved overall, we chose this buffer for our subsequent experiments.

### [Q3] How are community estimates affected by different lysis times and homogenisation?

After a short lysis time, i.e. 1h or 2h, we observed considerable variation among replicates in the community composition inferred. With an increasing duration of lysis, the relative abundance of a number of species was either consistently increasing or decreasing. For instance, in Experiment II, the relative abundance of *A. colemani* tended to decrease over time, while that of *D. yakuba* was increasing. Throughout lysis, the replicates remained visibly different from one another (Fig. 4).

Compared to mild lysis, homogenization substantially altered the community composition profiles inferred, with many species either increasing, or decreasing their relative abundance (Fig. 4). The smallest insects tended to be well detectable after mild lysis, whereas they yielded few or no reads after homogenization. For instance, the smallest species in our set (the parasitic wasp *Encarsia formosa*) was consistently represented by multiple reads after short lysis (1–4h) in Communities I and II (Experiment II). However, its proportion of reads decreased rapidly with increasing lysis time, and after homogenization, *E. formosa* was no longer detected in any replicate (Fig. 4a,b). The same was true for other small insects such as *Aphidius colemani, Aphidius ervi*, and *Aphidoletes aphidimyza* (Fig. 4). For bigger insects (e.g., the lepidopteran *Pieris napi*) and for harder-bodied insects (e.g., the mealworm beetle *Tenebrio molitor*), we observed the opposite tendency: species were detectable already after the shortest of lysis times (1h), but absolute read counts increased with lysis time and peaked after homogenization (Fig. 2). Species with hard exoskeleton, such as ants *Formica rufa* and *Formica fusca*, seem to have released little or no DNA in mild lysis, and thus homogenization was required to detect them (Fig. 4a-c).

Patterns revealed by nMDS ordination (Fig. 5) suggest different imprints of extraction method on different aspects of community composition. When comparing the composition of mock communities inferred from metabarcoding to their original, known composition, we noted that estimates derived from homogenates largely resemble the biomass composition of the community, while estimates from samples with shortest lysis times more closely resemble their original species abundance composition, with a clear trend as lysis time increases towards homogenization.

**Figure 5.**
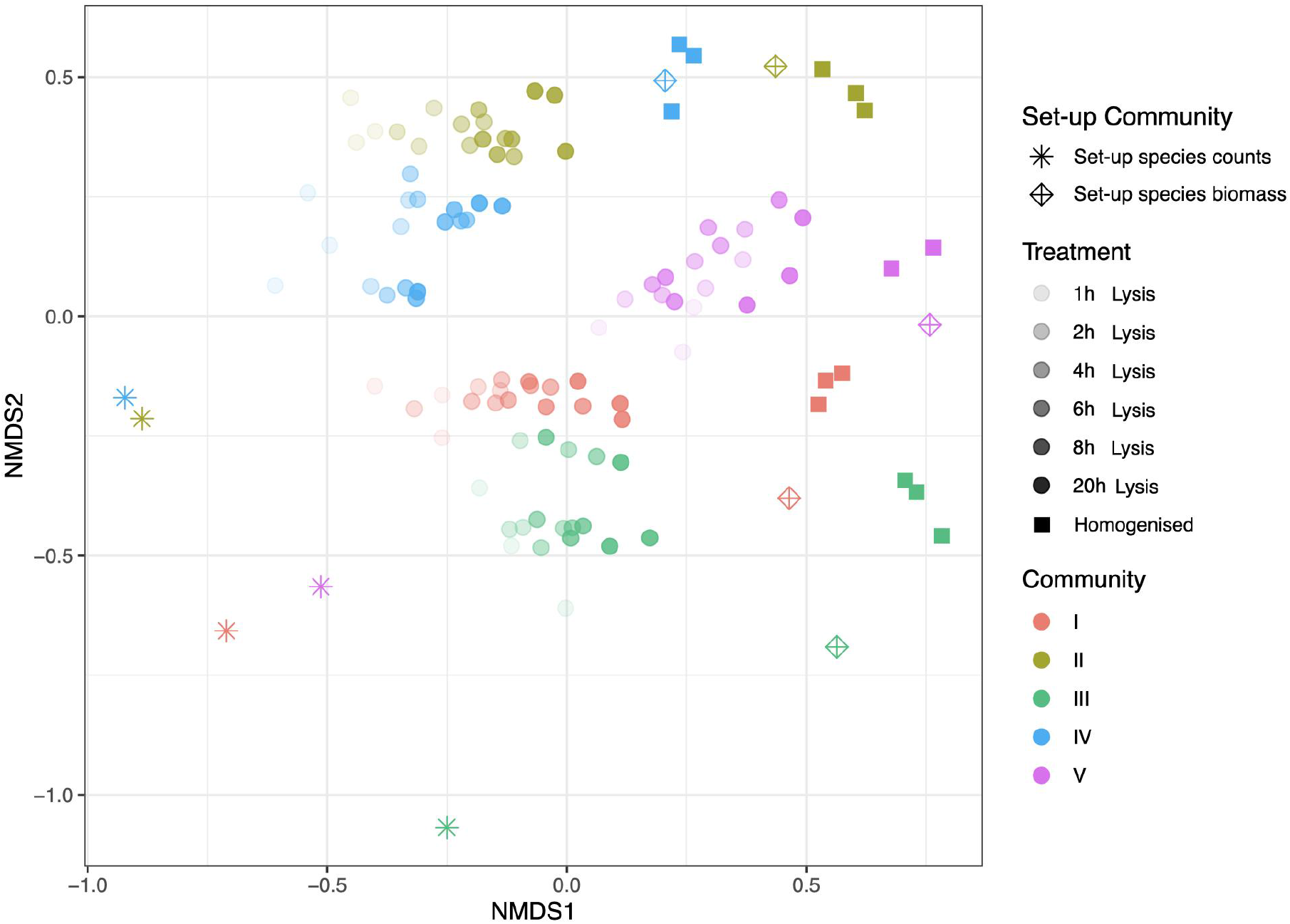
nMDS plot summing up COI reads for all samples in Experiment II. Five arthropod mock communities (I-V) in biological triplicates were subjected to mild lysis for 20h and sampled six times throughout the experiment and subsequently homogenized. Lysis times of each sample is represented in the graph as opacity level. Different shapes inform about whether a sample was homogenized (square), lysed (circle) or if it represents the original community set-up in terms of the relative number of individuals contributing to the community (asterisk) or the relative biomass of the species (crossed diamond).

### [Q4] Is it possible to obtain adequate species abundance estimates through the use of biological spike-ins?

Of the five spike-in species used in Experiment III, *Drosophila hydei* showed poor amplification and was therefore excluded from further comparisons. Thus, the following results apply to the four remaining spike-in species – *S. lateralis, G. bimaculatus, G. sigillatus and D. yakuba*.

The relative proportions of spike-in reads were similar both in the spike-in-only community, and the communities that included additional species, but there was considerable variation among replicates, especially in the mild lysis (Fig. S1). Using the probabilistic model for lysate data, there was strong evidence in favor of models accommodating species-specific differences in DNA yield over models that did not, while there was no evidence for such a difference in homogenates (Fig. 6). Posterior estimates of model parameters illustrate the same results in more detail (Figs 7, S2): there were distinct differences between species in average DNA yield per specimen in lysates (Fig. 7a) but not in homogenates (Fig. 7b).

**Figure 6.**
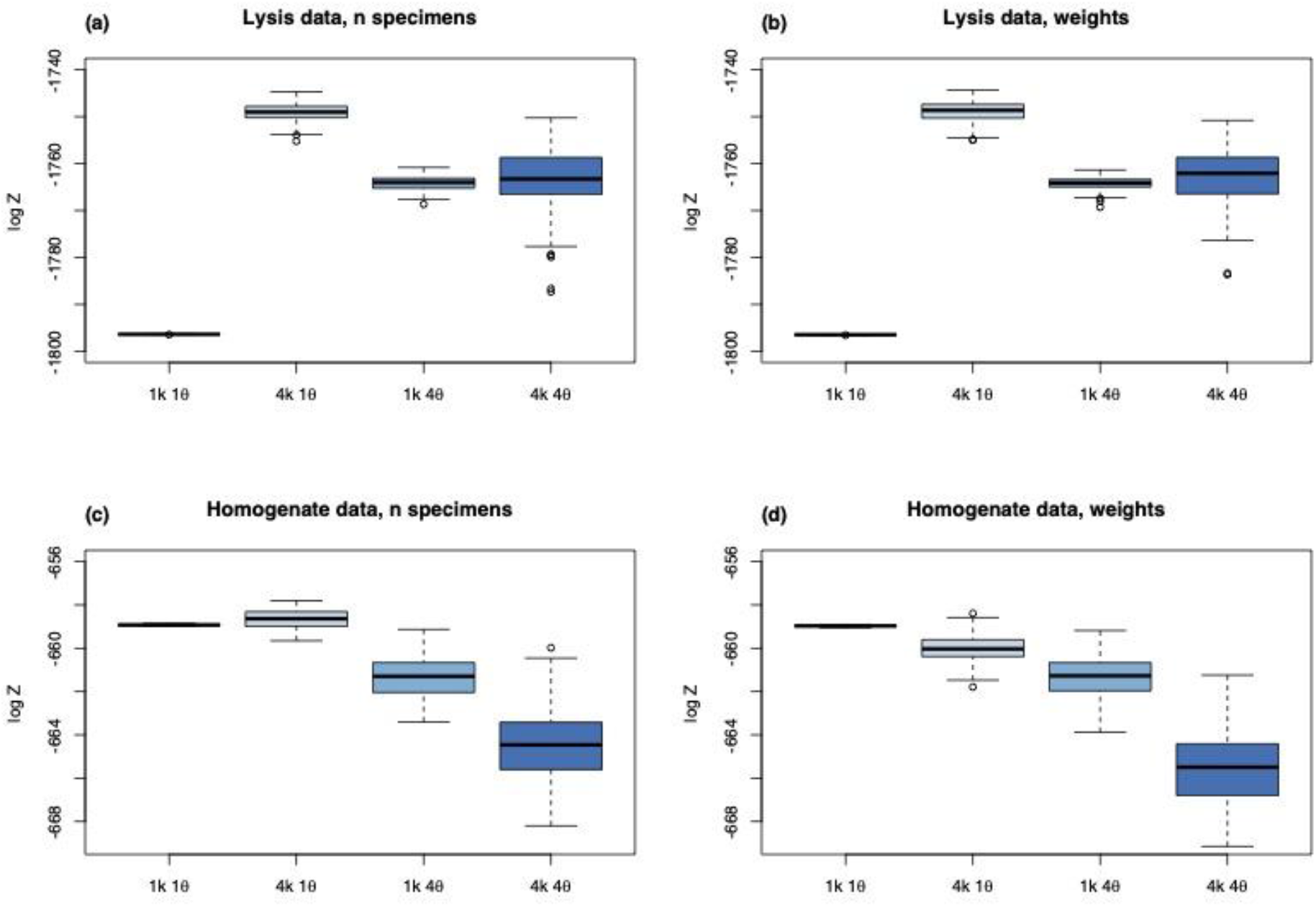
Estimated marginal likelihoods (log Z) for four models accommodating different sources of variation in the relative read numbers of the four spike-in species. Results are shown for lysis data in the top panels (a-b) and for homogenate data in the bottom panels (c-d). The left panels (a, c) show results for specimen composition and right panels (b,d) for biomass composition. A difference of five log likelihood units is considered very strong evidence against the model with lower likelihood (Kass & Raftery, 1995). In lysates, there is strong evidence for species-specific differences in DNA yield (4k over 1k models), whereas this is not the case in homogenates. In neither case, there is strong evidence for species-specific PCR amplification biases in these four species (4θ over 1θ models).

**Figure 7:**
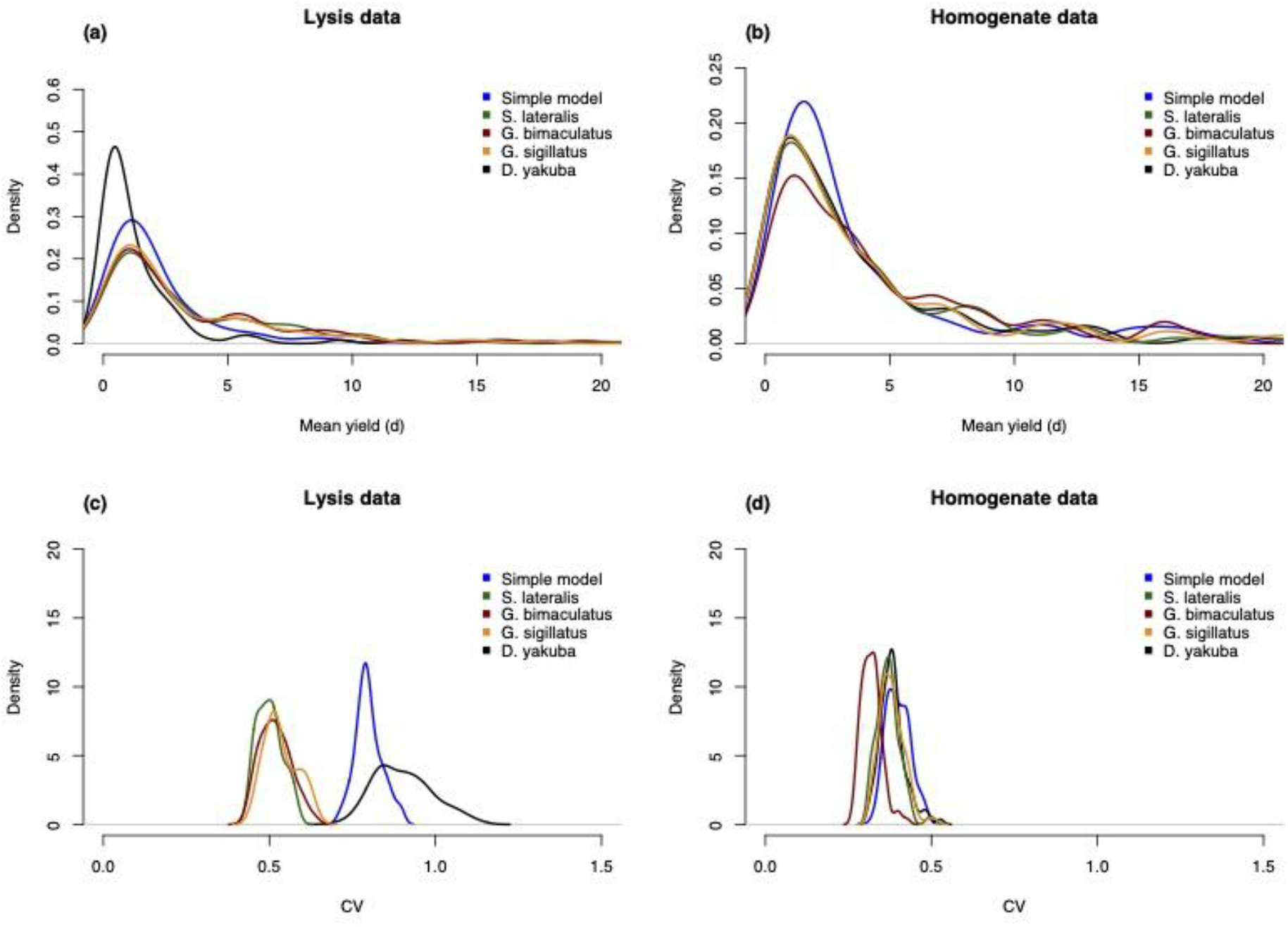
a-b shows the estimated posterior distributions of mean DNA yield per specimen, d, using two model variants: the “simple model” with no species-specific parameters (blue), and the model that allows the DNA yield of the four spike-in species to be different (remaining colors). In lysates (a), the mean DNA yield varies among the four species; in particular, the yield of Drosophila yakuba is lower than that of the other spike-in species. The DNA yield is more consistent across species in the homogenates (b). c-d shows the estimated posterior distribution for the coefficient of variance (CV) of DNA yield per specimen, using two model variants: the “simple model” with no species-specific parameters (blue), and the model that allows the DNA yield of the four spike-in species to be different (remaining colors). In lysates (c), the coefficient of variation was reduced to approximately 0.4 for three of the four species, but not for Drosophila yakuba. In homogenates (d), the coefficient of variation was estimated to be around 0.4 in the simple model, and accommodating species-specific DNA yields did not result in much improvement except for Gryllus bimaculatus.

Estimates of coefficients of variation can be used to pinpoint the level of consistency in read counts (the higher the CV, the lower consistency). Under the simple model that did not accommodate species-specific differences, the estimated CV for specimen numbers was around 0.8 for lysates (Fig. 7c) and around 0.45 for homogenates (Fig. 7d). In other words, the standard deviation (i.e., standard error, SE) was around 80% vs 45% of the mean, respectively. When species-specific differences in DNA yield were accommodated, the CV decreased to 0.5 for three of the four spike-ins in the lysate data, while it remained high for *Drosophila yakuba*, which was also the smallest of the four spike-ins. For the homogenate data, the CV estimates did not change when species-specific differences were accommodated.

When the spike-in data, fitted to a common model for all spike-in species, were used to estimate the abundances of the remaining species, we noticed clear species-specific effects: some species were consistently overestimated while others were consistently underestimated (Figs 8bd and S3bd). For lysate data, the estimates of specimen numbers were slightly better than the estimates of biomass (Figs 8a, S3a), while the opposite was true for homogenate data (Figs 8c, S3c). The lysate predictions tended to provide close to unbiased estimates of the number of specimens but clearly overestimate biomass, while the homogenate predictions gave approximately unbiased estimates of biomass but slightly underestimated the number of specimens.

**Figure 8:**
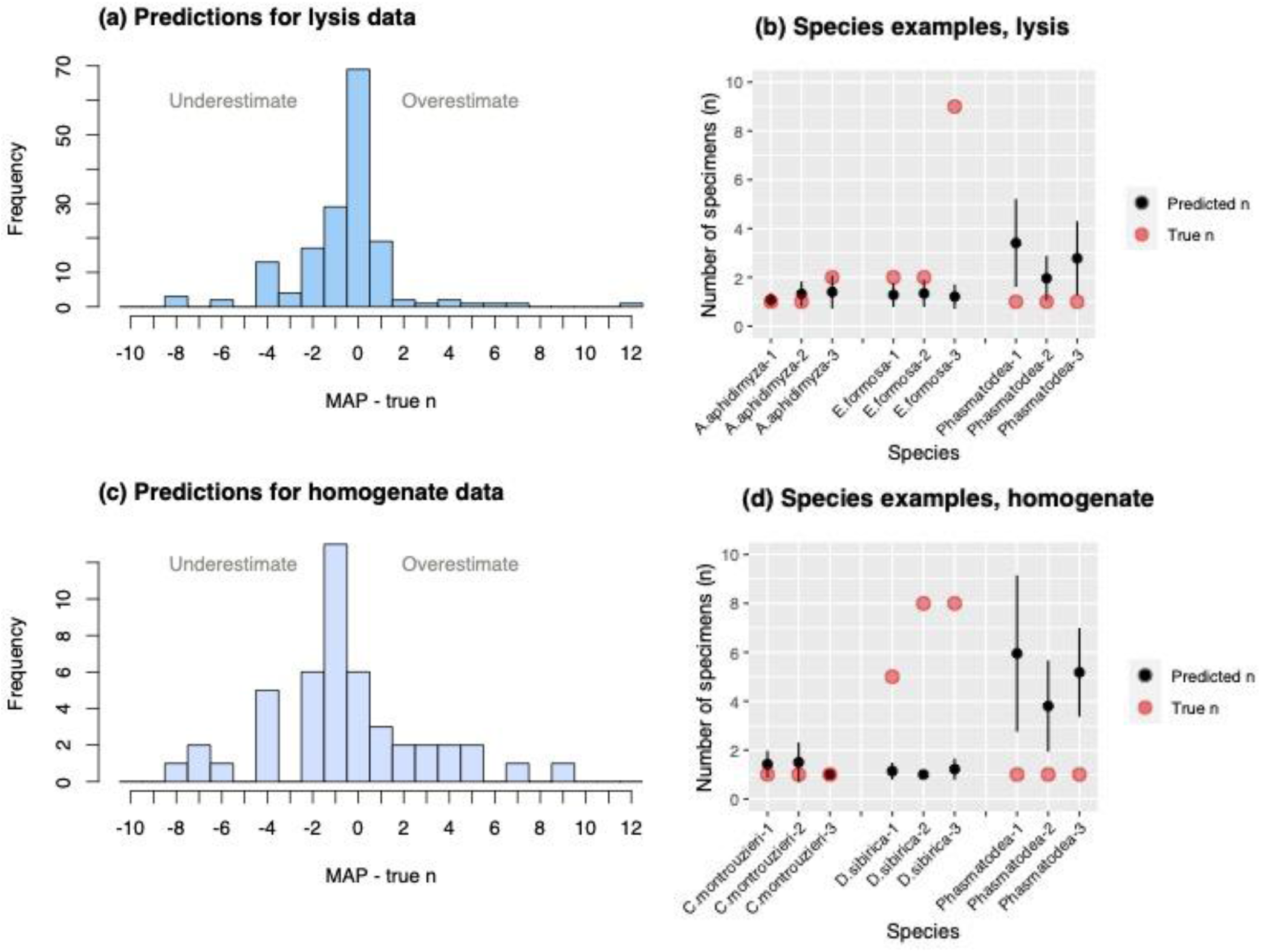
Predictions from Experiment III for the lysis treatment (a-b) and homogenate treatment (c-d). In (a) and (c) all predicted species and samples are combined. The predictions estimate how many specimens (n) for each of the seven predicted species are present in each sample. The histograms’ x-axes show the difference between the true value of n and the Maximum a posteriori (MAP) estimate. When the difference is zero, the model’s MAP estimate is also the true value of n. In (b) and (d), examples of species-specific predictions are shown: three samples each for three species, one species that is well predicted, one species that is underestimated and one species that is overestimated. The black circles mark the mean of the estimates, the bars are standard deviation, and the red circles mark the true value.

## Discussion

Our series of experiments provide significant new insights into methodological choices that determine the reliability of species detection, abundance and biomass estimations, in the metabarcoding of multi-species community samples. The use of replicate mock communities allowed us to assess the extent and significance of variation among seemingly identical insect assemblages. Using samples for the same mock communities that were collected at multiple timepoints during lysis as well as after homogenization, combined with biological spike-ins, we were able to assess the impact of selected methodological choices. Our results contribute to the understanding of trade-offs involved and the limits of DNA metabarcoding broadly. Below, we explore different aspects of the results.

### Reproducibility of metabarcoding results

Previous studies have reported limited differences in the perception of community composition gained from parallel technical replicates. Such findings were made by, e.g., Marquina et al. (2019) and Nielsen et al. (2019) when comparing different methods of DNA purification using lysate from the same bulk sample. Reproducibility was also evaluated by Martoni and collaborators (2022), who used mock communities to evaluate taxonomic bias introduced by the choice of PCR primers and by the library preparation method. However, to our knowledge, the current study is the first to examine variation between biological replicates of multi-species communities.

Our results reveal a surprising amount of variation among identical replicates of the same mock community. This variation concerns both relative read abundances and the presence/absence of species. It was particularly strong after mild lysis treatment but also persisted, to a lesser extent, after homogenisation. In fact, samples representing different replicates of the same mock community were occasionally less similar to each other than to samples of distinct mock communities.

It seems likely that the lack of reproducibility is largely due to natural variation in DNA yield among individuals of the same species. Recall that in our study, efforts were made to limit variation among individuals; for instance, we used only one life stage per species, and excluded individuals that appeared poorly sclerotized, injured, particularly large or small, or otherwise unusual. Thus, one might expect even more variation among real wild-caught insects, where specimens differ in life stage, age, sex, health, and physiological state. Adding to this, how samples are handled, transported, and stored, will also contribute to variation in DNA yield of specimens, with the effects likely to vary among species (Marquina et al., 2021). Thus, the variation is inherent in metabarcoding procedures and will clearly limit the precision of any estimates of community composition.

### Trade-offs between protocols

The different outcomes of our metabarcoding experiments contribute to the understanding of the trade-offs involved. Generally speaking, a milder protocol (short lysis and mild buffer) results in more accurate reflection of the presence/absence of species, and community profiles more closely resembling those based on numbers of specimens per species. The more aggressive the protocol (i.e., buffer with stronger lytic properties, longer lysis, homogenization), the more reflective the results are of the biomass composition of the community. This pattern is perhaps most clearly seen in the nMDS plot from Experiment II, where the shortest lysis time (1h) results in estimates relatively close to the specimen composition of the community. As lysis time increases, the results move towards the biomass composition of the community and homogenates represent the extreme endpoint of this trend.

The comparison of different buffers in Experiment I can be interpreted in similar terms. The milder Buffer 1 yields relatively good coverage of a broad range of insect species while the more aggressive Buffer 2 yields more sequences of larger species, at the expense of the representation of small insects. Notably, the volume of lysate and homogenate aliquots used for DNA extraction was the same across all treatments and the libraries were prepared the same way, although the input DNA concentration and read numbers may have differed. Thus, the shifting representation of species in metabarcoding is reflecting the varying proportion of their DNA template in the starting aliquot. The likely explanation for this is that small insects have a high surface-to-volume ratio and tend to release relatively more DNA initially. With more aggressive treatments the DNA pool becomes more reflective of the insects’ volumes. As a result, large species dominate in terms of available DNA and we fail to detect small species. This is demonstrated by the high rate of false negatives for some of the smallest species (*Aphidoletes aphidimyza, Dacnusa sibirica, Encarsia formosa* and *Tuberculatus annulatus*) in the homogenates of Experiment II.

On top of this, there are clear species-specific effects, the most obvious of which is the degree of sclerotization. For instance, *Hermetia illucens* larvae yielded little DNA in mild lysis treatments despite their large size. This may be due to their unusually hard integument (Ståhls et al., 2020). Similarly, in Experiment II, the two ants (*Formica fusca* and *Formica rufa*) often required homogenization to release enough DNA for detection. Still, their low representation even in homogenates suggests that this effect might be enhanced by PCR bias against them. Undoubtedly, there are also other traits that influence how species react to different extraction protocols.

Overall, our findings are in line with the results of Marquina et al. (2022), who found that small and soft species were more difficult to detect after treatment with a chemically more aggressive buffer (higher SDS, DTT, and proteinase K concentrations) than after lysis with the milder buffer corresponding to Buffer 1 in the current study. They also agree with the results of Marquina et al. (2019), who showed that hard-bodied and large insects are more easily detected in homogenates, while analysis of preservative ethanol – similar to extremely mild lysis – is better for the detection of small and soft-bodied insects. However, it remains to be evaluated whether and under what conditions this pattern holds true for real terrestrial arthropod catches (but see Nielsen et al. 2019).

### Abundance estimates from metabarcoding data

How well species abundances may be estimated through metabarcoding is currently a hotly debated topic. Early studies suggested that metabarcoding was unsuitable for quantification, but more recent efforts using correction factors or spike-ins of known concentration to calibrate metabarcoding-based estimates have reported encouraging results (Ershova et al., 2021; Krehenwinkel et al., 2017). Nevertheless, our study offers some of the first quantitative analyses of the accuracy that might be achieved using such approaches.

The use of biological spike-ins is based on the expectation that parameters learned from spike-in data can be applied to the species occurring naturally in the sample, but several sources of variation may influence the results. First, the mean DNA yield per specimen is likely to differ dramatically among species. Second, specimens from a single species can vary substantially in DNA yield. Finally, PCR amplification may also vary considerably among species. Indeed, in Experiment III, mean DNA yields varied considerably across the spike-ins in lysates, making it difficult to calibrate data for species other than those deliberately spiked-in. However, if species-specific calibration data are available, it seems possible to bring down the error of abundance estimates for specimens to a CV of around 50% of the mean under ideal conditions (recall that we size-standardized the spike-in specimens). The mean DNA yield in the homogenate data is more uniform across species, and the CV estimates are lower. Under ideal conditions, one might then be able to reach a CV of 40% when using homogenate data for spike-in species to calibrate biomass estimates for other species. These results, however, do not apply to species with strong PCR amplification biases such as *Drosophila hydei* and the stick insect (Phasmatodea) - with very low and high PCR factors, respectively.

Overall, our data suggest that there is a natural limit to the precision of metabarcoding-based abundance estimates. Specifically, it appears difficult to bring the standard error of homogenate-based estimates much below 40-45% of the mean when using biological spike-in references. These limits derive from natural variation between specimens, and stochastic processes in sample preparation. Nonetheless, the statistical method presented here can still be improved to increase the range of situations in which this natural limit can be reached. For instance, the error associated with variable DNA yield from biological spike-ins could possibly be reduced through synthetic spike-ins, such as linearized plasmids with artificial COI targets that are added in pre-defined quantities to homogenates (Palmer et al., 2018; Tourlousse et al., 2017). Alternatively, using individual-specimen barcoding of a subset of lysed or homogenized samples could extend the calibration data to species that occur naturally in the samples. Such an approach would also open up new possibilities for model-free methods by providing a sufficiently large training dataset for machine learning algorithms.

### Towards the optimal protocol

Given the different advantages and disadvantages of lysis versus homogenization, it will be hard to strike the ideal compromise. To find the smallest and rarest species in the sample will always be difficult – and particularly difficult for homogenate data, where they constitute a minuscule proportion of the total DNA. One approach, albeit destructive and costly, might be metabarcoding of both mild-lysis and homogenates of the same sample. Such a combination would take advantage of the combined qualitative and quantitative strong suits of the two approaches.

With respect to the specifics of the mild lysis protocol, our results suggest that intermediate lysis times using our preferred buffer (Buffer 1) give robust results, and that community profiles obtained after 2h digestion are comparable to those after 4h, 6h, or 8h. Digestion for 1h appears to be suboptimal, with increased variation in species representation and lower overall DNA yield. Samples digested for 20h start to resemble homogenates, with decreased representation of small species.

All in all, our current analysis provides key pointers for the design of an optimal protocol for processing bulk insect samples, with the specifics to be further attuned to local logistics and laboratory setup. We remain optimistic about the prospects of improved statistical modeling to unlock the resulting data’s full potential.

## Author Contributions

E.I., P.L. and F.R. conceived the ideas and designed methodology; E.I. and P.L. collected the data with the assistance of M.B. and M.P.; E.I. and E.G. analyzed the data; E.I., P.L. and F.R. led the writing of the manuscript. All authors contributed critically to the drafts and gave final approval for publication.

All authors declare that they have no conflicts of interest.

## Data Availability

Raw data will be made available on The European Nucleotide Archive (ENA) at the time of publication.

## Acknowledgements

We thank Jan Kudlicka at BI Norwegian Business School for assistance with Birch and model discussions.This project was supported by the Knut and Alice Wallenberg Foundation (KAW 2017.088 to FR), Swedish Research Council grant No 2018-04620 (to FR), Polish National Agency for Academic Exchange grant PPN/PPO/2018/1/00015 (to PL), Polish National Science Centre grant 2018/31/B/NZ8/01158 (to PL) and the National Institute Of General Medical Sciences of the NIH under award number R35GM124701 to Brandon S. Cooper (insect culture). TR was funded by the European Research Council Synergy Grant 856506 (LIFEPLAN) and a Career Support grant from the Swedish University of Agricultural Sciences. AJMT was funded by the Swedish Research Council (2019-04493). The data handling was enabled by resources provided by the Swedish National Infrastructure for Computing (SNIC), partially funded by the Swedish Research Council through grant agreement no. 2018-05973.

