## Supplementary information for "Optimizing insect metabarcoding using replicated mock communities"

### Supporting Information for the "Optimizing insect metabarcoding using replicated mock communities" paper:

**Text S1. Experiment I: Assessing the effects of buffer and lysis time/homogenization**

**Text S2. Library preparation details**

**Fig. S1. Variance and standard deviation in relative count numbers for each spike-in species**

**Fig. S2. Estimated posterior distributions of mean DNA yield per biomass**

**Fig. S3. Estimated posterior distribution for the coefficient of variance (CV) of DNA yield per biomass**

**Fig. S4. Predictions from Experiments_III, for the mild lysis and homogenate treatment**

**Table S1. Summary of reference insect species**

**Table S2. Overview of species used in the mock communities in Experiment I**

**Table S3. Overview of species used in the mock communities in Experiment II**

**Table S4. Overview of species used in the mock communities in Experiment III**

**Table S5. Summary of COI sequencing results - numbers of reads per Experiment**

### TEXT S1: Experiment I: Assessing the effects of buffer and lysis time/homogenization (Q2, Q3)

To look into the effect of different lysis buffers we used 13 species to assemble three mock communities: community "S" consisted of 11 insects belonging to 8 species; "M" had 19 individuals of 12 species; and "L" had 25 insects of 13 species representing different morphologies (Fig.1, Table S2). Each mock community was prepared in six replicates: three were subjected to lysis with Buffer 1 (same as used in later experiments) while the other three were lysed with Buffer 2.

Buffer 1 was prepared according to Vesterinen et al. (2016; modified from Aljanabi and Martinez (1997)). It consisted of 0.4M NaCl, 10mM Tris-HCl pH=8.0, 2mM EDTA pH=8, 2% SDS, 0,1% proteinase K (20mg/mL) and molecular biology grade water.

Buffer 2 was prepared according to a recipe for arthropod specific lysis buffer from the Canadian Centre for DNA Barcoding (CCDB; Ivanova et al., 2006). It consisted of 700mM GuSCN, 30mM EDTA, 30mM Tris-HCl pH=8, 0.5% Triton X-100, 5% Tween-20.

Samples were incubated in 7mL tubes in 5mL of the lysis buffer, in a shaking water bath at 56°C. 50uL aliquots were taken after 2h, 4h and 6h of incubation. Afterwards, the samples and the remaining lysate were homogenised using Bead Ruptor Elite (OMNI INTERNATIONAL). DNA was purified from 20µL aliquots of the lysate or homogenate using 30µL of Carboxylate-modified Sera-Mag SpeedBeads (Cytiva) in a PEG/NaCl buffer, and eluted in 20µL 1x TE buffer (protocol at:

https://protocols.io/view/pcr-cleaning-using-home-made-spri-magnetic-beads-m-b2gdqbs6)
.

### TEXT S2: Library preparation

We prepared COI amplicon libraries using a two-step PCR approach, as described in Marquina et al. (2021). In short, in PCR1, we amplified the 458-bp target region of the COI gene using primer pair BF3-BR2 (Elbrecht et al., 2019) with variable-length inserts and Illumina adapter tails. The thermocycling protocol included an activation phase at 95°C for 15min followed by 25 cycles of 30s denaturalization at 94°C, 90s of annealing at 50°C and 90s of extension at 72°C, followed by the final extension at 72°C for 10min. Subsequently, the product was cleaned using Sera-Mag SpeedBeads magnetic bead solution, and eluted in 10μL of TEx1 buffer, of which 1μL was used as template for the indexing PCR2, which added Illumina adapters with barcode combinations unique for each sample (sequences in Table S11). PCR2 conditions were the same as PCR1, except that only 5 cycles were used. Subsamples of products were checked on 2.5% agarose gel and pooled equimolarly based on band brightness. Pools were cleaned again with Sera-Mag SpeedBeads and sequenced, alongside samples from other projects, on three Illumina MiSeq v3 lanes (2×300bp reads) at the Institute of Environmental Sciences of Jagiellonian University (Kraków, Poland).

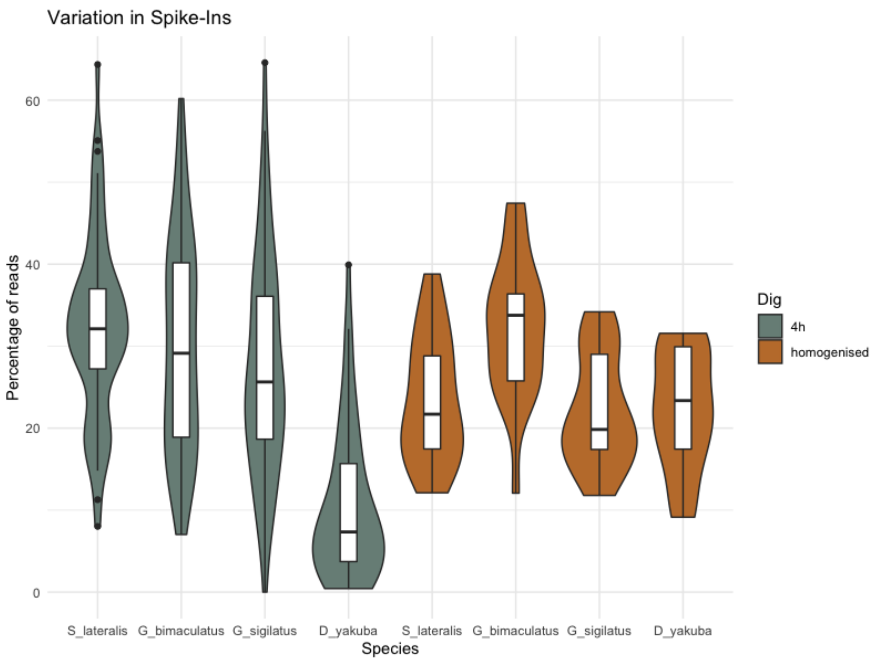


***Figure S1. Variance and standard deviation in relative count numbers for each spike-in species*** *in Experiment III. Plot shows the extent of variation in read counts after 4 hours of mild lysis (grey) (n=58) and after homogenisation (mustard) (n=22).*

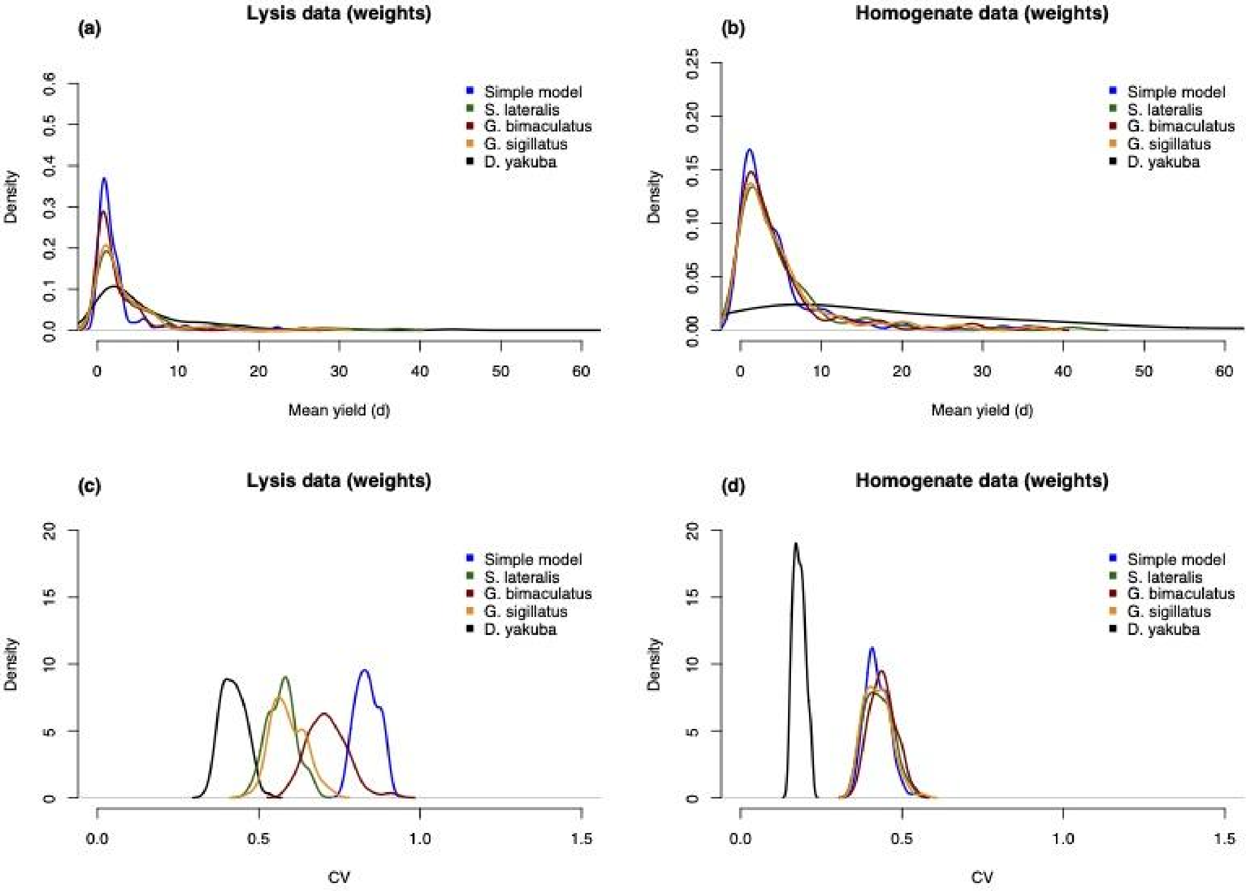


***Figure S2 a-b: Estimated posterior distributions of mean DNA yield per biomass**, d, using two model variants: the "simple model" with no species-specific parameters (blue), and the model that allows the DNA yield of the four spike-in species to be different (remaining colors). In lysates (a), the mean DNA yield varies among the four species, although the differences are not as clear as for DNA yield per specimen (Fig. 8). Here, the yield per mass of* Drosophila yakuba *is higher than that of the other spike-in species and some variation can be seen for the remaining three species. The DNA yield per biomass is very similar for three species, while there is considerable uncertainty concerning the DNA yield for* Drosophila yakuba, *resulting in the absence of strong evidence that the yield differs from that of the other species (b). The latter result could potentially be due to the inaccuracies we introduced by using a standard weight for each species rather than weighing each sequenced specimen. **c-d: Estimated posterior distribution for the coefficient of variance (CV) of DNA yield per biomass**, using two model variants: the "simple model" with no species-specific parameters (blue), and the model that allows the DNA yield of the four spike-in species to be different (remaining colors). In lysates (c), the coefficient of variation was reduced to different degrees across the four spike-in species, but only for* Drosophila yakuba *to values below 0.5. In homogenates (d), the coefficient of variation was estimated to be around 0.4 in the simple model. Assuming species-specific DNA yields improved the coefficient of variation considerably for* Drosophila yakuba*, but had negligible effect on the other species.*

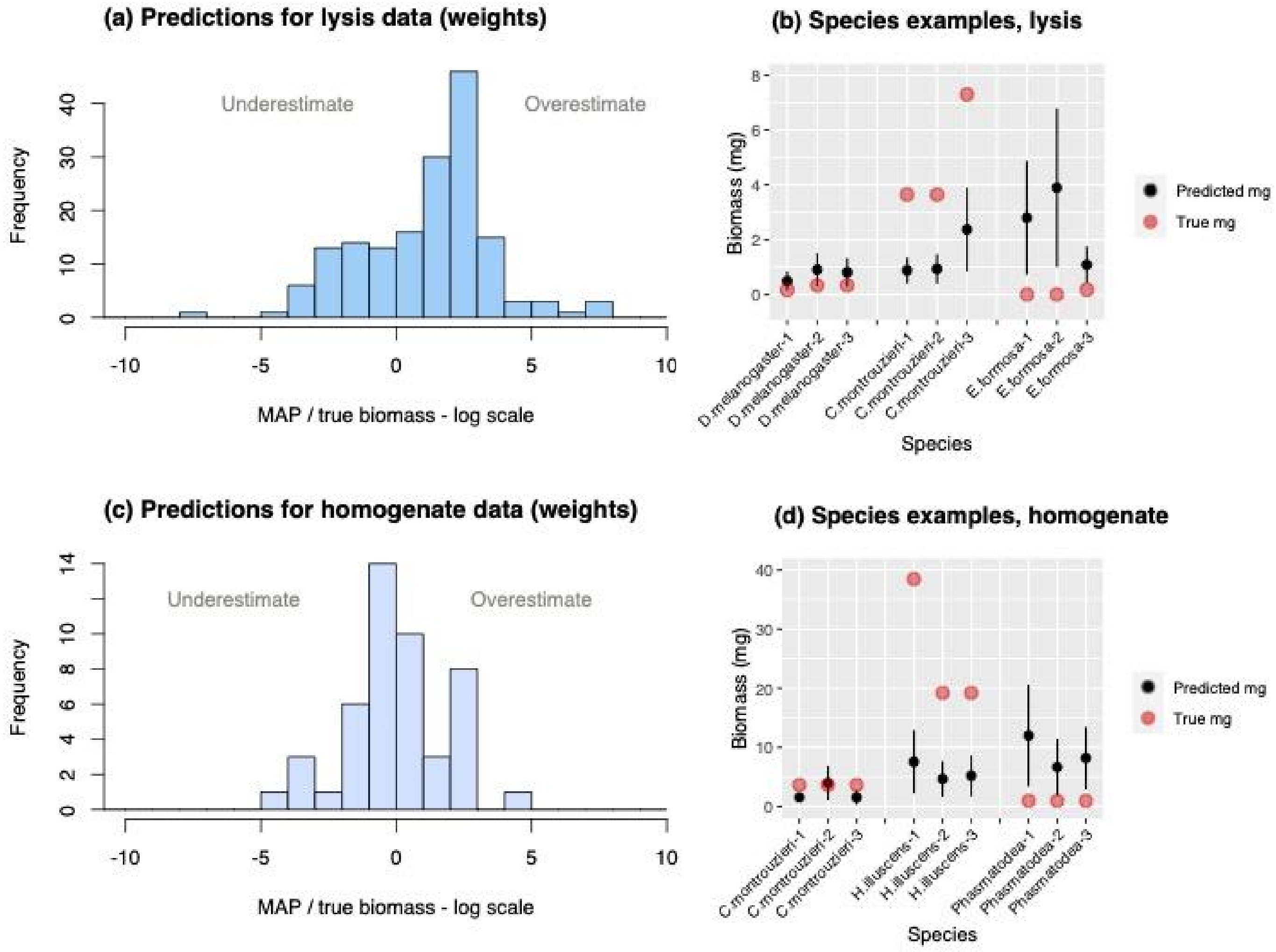


***Figure S3: Predictions from Experiments III, for the lysis treatment (a-b) and homogenate treatment (c-d).*** *In (a) and (c) all predicted species and samples are combined. The predictions estimate how much biomass (mg) for each of the seven predicted species are present in each sample. The plots' x-axes show MAP/true biomass on a log scale. At zero on the x axis, the model's MAP estimate is also the true biomass value. In (b) and (d) examples of species-specific predictions are shown: three samples each for three species, one species that is well predicted, one species that is underestimated and one species that is overestimated. The black circles mark the mean of the estimates, the bars are standard deviation, and the red circles mark the true value.*

***Table S1. Summary of reference insect species.*** *We constructed communities using standardized, reference insect species. We obtained a total of 31 species from different sources, including biocontrol stores, pet stores, hobbyists and laboratory cultures. All insects were preserved in 90% ethanol, and stored at -20°C until used. If the initial material consisted of juveniles at different life stages, we standardized their sizes by selecting one developmental stage. We obtained dry weights for each species by weighting a set of ten representative specimens and calculating an average individual weight. Mock communities were prepared by combining pre-set numbers of individuals from each species in a Falcon tube with ethanol, which was then drained prior to lysis. Every mock community was prepared in three identical copies. The composition of mock communities varied across experiments, and each experiment used different community types, as explained below; see Tables S2-4 for details.*

| | | | | | **Used in Exp.** | | | | |
|---|---|---|---|---|---|---|---|---|---|
| **#** | **Species** | **Order** | **Family** | **Developmental stage** | **I** | **II** | **III** | **Dry weight [mg] per indiv.** | **Origin** |
| 1 | ***Acheta domesticus*** | Orthoptera | Gryllidae | size-selected nymphs | NO | YES | NO | 3.55 | commercial culture (pet food) - reptilgrottan.se |
| 2 | ***Aphidius colemani*** | Hymenoptera | Braconidae | newly eclosed adults | YES | YES | NO | 0.04 | commercial culture (biocontrol) - koppert.se |
| 3 | ***Aphidius ervi*** | Hymenoptera | Braconidae | newly eclosed adults | NO | YES | NO | 0.09 | commercial culture (biocontrol) - koppert.se |
| 4 | ***Aphidoletes aphidimyza*** | Diptera | Cecidomyiida e | newly eclosed adults | NO | YES | YES | 0.07 | commercial culture (biocontrol) - koppert.se |
| 5 | ***Blaptica dubia*** | Blattodea | Blaberidae | size-selected nymphs | NO | YES | NO | 65.66 | commercial culture (pet food) - reptilgrottan.se |
| 6 | ***Calliphora vomitoria*** | Diptera | Calliphoridae | newly eclosed adults | NO | YES | NO | 14.35 | commercial culture (fishing store) - Fibe AB, Sweden |
| 7 | ***Cryptolaemus montrouzieri*** | Coleoptera | Coccinellidae | adults | YES | NO | YES | 3.65 | commercial culture (biocontrol) - koppert.se |
| 8 | ***Dacnusa sibirica*** | Hymenoptera | Braconidae | adults | YES | YES | YES | 0.09 | commercial culture (biocontrol) - koppert.se |
| 9 | ***Dermestes haemorrhoid alis*** | Coleoptera | Dermestidae | adults | NO | YES | NO | 14.95 | dermestarium, Swedish Museum of Natural History |
| 10 | ***Drosophila hydei*** | Diptera | Drosophilidae | adults | YES | NO | YES | 0.42 | commercial culture (pet food) - reptilgrottan.se |
| 11 | ***Drosophila melanogaster*** | Diptera | Drosophilidae | adults | YES | NO | YES | 0.17 | lab culture - Brandon Cooper, University of Montana |
| 12 | ***Drosophila simulans*** | Diptera | Drosophilidae | adults | NO | YES | NO | 0.35 | lab culture - Brandon Cooper, University of Montana |
| 13 | ***Drosophila teissieri*** | Diptera | Drosophilidae | adults | NO | YES | NO | 0.23 | lab culture - Brandon Cooper, University of Montana |
| 14 | ***Drosophila yakuba*** | Diptera | Drosophilidae | adults | YES | YES | YES | 0.21 | lab culture - Brandon Cooper, University of Montana |
| 15 | ***Encarsia formosa*** | Hymenoptera | Aphelinidae | newly eclosed adults | YES | YES | YES | 0.0015 | commercial culture (biocontrol) - koppert.se |

| | | | | | | | | | |
|---|---|---|---|---|---|---|---|---|---|
| **16** | ***Formica fusca*** | Hymenoptera | Formicidae | adult workers | NO | YES | NO | 1.83 | wild colony, collected near Swedish Museum of Natural History |
| **17** | ***Formica rufa*** | Hymenoptera | Formicidae | adult workers | NO | YES | NO | 2.62 | wild colony, collected near Swedish Museum of Natural History |
| **18** | ***Gryllodes sigillatus*** | Orthoptera | Gryllidae | size-selected nymphs | YES | NO | YES | 1.2 | commercial culture - cricketsfarm.pl, Poland |
| **19** | ***Gryllus bimaculatus*** | Orthoptera | Gryllidae | size-selected nymphs | YES | NO | YES | 1.81 | commercial culture - e-robaczek.pl, Poland |
| **20** | ***Hermetia illucens*** | Diptera | Stratiomyidae | size-selected larvae | YES | NO | YES | 19.24 | commercial culture - egzotic-room.com, Poland |
| **21** | ***Lasiommata megera*** | Lepidoptera | Nymphalidae | size-selected larvae | NO | YES | NO | 5.17 | lab culture - Mats Ittonen, Stockholm U |
| **22** | ***Locusta migratoria*** | Orthoptera | Acrididae | size-selected nymphs | NO | YES | NO | 29.21 | commercial culture (pet food) - reptilgrottan.se |
| **23** | ***Macrolophus pygmaeus*** | Hemiptera | Miridae | adults | YES | YES | NO | 0.5 | commercial culture (biocontrol) - koppert.se |
| **24** | ***Necrobia rufipes*** | Coleoptera | Cleridae | adults | NO | YES | NO | 4.51 | dermestarium, Swedish Museum of Natural History |
| **25** | ***Pieris napi*** | Lepidoptera | Pieridae | adults | NO | YES | NO | 21.45 | lab culture - Christer Wiklund Stockholm U |
| **26** | ***Polygonia c-album*** | Lepidoptera | Nymphalidae | adults | NO | YES | NO | 38.8 | lab culture - Maertha Eriksson, Stockholm U |
| **27** | ***Shelfordella lateralis*** | Blattodea | Blattidae | size-selected nymphs | YES | NO | YES | 1.27 | commercial culture - e-robaczek.pl, Poland |
| **28** | **Stick insect** | Phasmatodea | | newly hatched nymphs | YES | YES | YES | 0.98 | lab culture - Niklas Apelqvist, Swedish Museum of Natural History |
| **29** | ***Tenebrio molitor*** | Coleoptera | Tenebrionida e | adults | NO | YES | NO | 50.79 | lab culture - Anna Michalik, Jagiellonian U |
| **30** | ***Thermobia domestica*** | Zygentoma | Lepismatidae | adults | NO | YES | NO | 30 | lab culture - Anna Michalik, Jagiellonian U |
| **31** | ***Tuberculatus annulatus*** | Hemiptera | Aphididae | adults | NO | YES | NO | 0.05 | lab culture - Laura van Dijk, Stockholm U |

***Table S2. Experimental design of Experiment I.*** *Numbers of individuals, per species, used to make mock communities.*

| No. | Species | Order | Comm. S | Comm. M | Comm. L | Total no. used |
|---|---|---|---|---|---|---|
| 1 | *Gryllus bimaculatus* | Orthoptera | 1 | 1 | 1 | 18 |
| 2 | *Shelfordella lateralis* | Blattodea | 1 | 1 | 1 | 18 |
| 3 | *Gryllodes sigillatus* | Orthoptera | 1 | 1 | 1 | 18 |
| 4 | *Drosophila yakuba* | Diptera | 1 | 1 | 1 | 18 |
| 5 | *Drosophila hydei* | Diptera | 1 | 1 | 1 | 18 |
| 6 | *Encarsia formosa* | Hymenoptera | 0 | 1 | 1 | 12 |
| 7 | *Drosophila melanogaster* | Diptera | 2 | 5 | 3 | 60 |
| 8 | *Dacnusa sibirica* | Hymenoptera | 3 | 1 | 5 | 54 |
| 9 | *Cryptolaemus montrouzieri* | Coleoptera | 0 | 1 | 3 | 24 |
| 10 | stick insect | Phasmatodea | 0 | 0 | 1 | 6 |
| 11 | *Hermetia illucens* | Diptera | 1 | 1 | 2 | 24 |
| 12 | *Macrolophus pygmaeus* | Hemiptera | 0 | 2 | 3 | 30 |
| 13 | *Aphidius colemani* | Hymenoptera | 0 | 3 | 2 | 30 |
| Replicates | | | 6 | 6 | 6 | |
| Total number of insects per sample | | | 11 | 19 | 25 | |

***Table S3. Experimental design of Experiment II.*** *Numbers of individuals and approximated biomass, per species, used to make mock communities in Experiment II.*

| | | Community 1 | | Community 2 | | Community 3 | | Community 4 | | Community 5 | |
|---|---|---|---|---|---|---|---|---|---|---|---|
| | **Species** | # individuals | weight [mg] | # individuals | weight [mg] | # individuals | weight [mg] | # individuals | weight [mg] | # individuals | weight [mg] |
| **1** | *Acheta domesticus* | 1 | 3.55 | 5 | 17.76 | 1 | 3.55 | 1 | 3.55 | 5 | 17.76 |
| **2** | *Aphidius colemani* | 5 | 0.21 | 10 | 0.43 | 1 | 0.04 | 30 | 1.28 | 1 | 0.04 |
| **3** | *Aphidius ervi* | 5 | 0.46 | 1 | 0.09 | 5 | 0.46 | 1 | 0.09 | 0 | 0 |
| **4** | *Aphidoletes aphidimyza* | 5 | 0.35 | 0 | 0 | 10 | 0.7 | 1 | 0.07 | 3 | 0.21 |
| **5** | *Blaptica dubia* | 1 | 65.66 | 0 | 0 | 1 | 65.66 | 1 | 65.66 | 3 | 196.98 |
| **6** | *Calliphora vomitoria* | 1 | 14.35 | 1 | 14.35 | 1 | 14.35 | 3 | 43.05 | 3 | 43.05 |
| **7** | *Dacnusa sibirica* | 5 | 0.44 | 10 | 0.88 | 1 | 0.09 | 30 | 2.63 | 5 | 0.44 |
| **8** | *Dermestes haemorrhoidalis* | 1 | 14.95 | 1 | 14.95 | 1 | 14.95 | 0 | 0 | 1 | 14.95 |
| **9** | *D. melanogaster* | 10 | 1.69 | 1 | 0.17 | 10 | 1.69 | 1 | 0.17 | 30 | 5.07 |
| **10** | *D. simulans* | 10 | 3.49 | 10 | 3.49 | 1 | 0.35 | 30 | 10.46 | 1 | 0.35 |
| **11** | *D. yakuba* | 20 | 4.18 | 13 | 2.72 | 1 | 0.21 | 6 | 1.26 | 11 | 2.3 |

| | | | | | | | | | | | |
|---|---|---|---|---|---|---|---|---|---|---|---|
| **12** | *D. teissieri* | 10 | 2.34 | 1 | 0.23 | 10 | 2.34 | 1 | 0.23 | 0 | 0 |
| **13** | *Encarsia formosa* | 30 | 0.045 | 30 | 0.045 | 1 | 0.0015 | 10 | 0.015 | 10 | 0.015 |
| **14** | *Formica fusca* | 1 | 1.83 | 3 | 5.5 | 1 | 1.83 | 5 | 9.16 | 0 | 0 |
| **15** | *Formica rufa* | 1 | 2.62 | 1 | 2.62 | 3 | 7.86 | 0 | 0 | 3 | 7.86 |
| **16** | *Lassiomata megera* | 1 | 5.17 | 1 | 5.17 | 0 | 0 | 1 | 5.17 | 1 | 5.17 |
| **17** | *Locusta migratoria* | 1 | 29.21 | 1 | 29.21 | 3 | 87.64 | 0 | 0 | 1 | 29.21 |
| **18** | *Macrolophus pygmaeus* | 5 | 2.5 | 10 | 5 | 1 | 0.5 | 1 | 0.5 | 0 | 0 |
| **19** | *Necrobia rufipes* | 1 | 4.51 | 1 | 4.51 | 3 | 13.54 | 0 | 0 | 1 | 4.51 |
| **20** | *Pieris napi* | 1 | 21.45 | 0 | 0 | 1 | 21.45 | 1 | 21.45 | 0 | 0 |
| **21** | *Polygonia c album* | 1 | 38.8 | 3 | 116.39 | 1 | 38.8 | 1 | 38.8 | 1 | 38.8 |
| **22** | Phasmatodea | 1 | 0.98 | 1 | 0.98 | 0 | 0 | 1 | 0.98 | 3 | 2.94 |
| **23** | *Tenebrio molitor* | 1 | 50.79 | 1 | 50.79 | 0 | 0 | 1 | 50.79 | 2 | 101.57 |
| **24** | *Thermobia domestica* | 1 | 30 | 0 | 0 | 1 | 30 | 1 | 30 | 3 | 90 |
| **25** | *Tuberculatus annulatus* | 5 | 0.25 | 1 | 0.05 | 10 | 0.5 | 5 | 0.25 | 3 | 0.15 |
| | **Sum of individuals:** | 124 | | 106 | | 68 | | 132 | | 91 | |

***Table S4. Experimental design for Experiment III.*** *Numbers of individuals, per species, used to make mock communities in Experiment III.*

| **Comm** | *S. lateralis* | *G. bimaculatus* | *G. sigillatus* | *D. yakuba* | *D. hydei* | *H. illucens* | *Stick insect* | *C. mantrouzieri* | *D. sibirica* | *D. melanogaster* | *A. aphidimyza* | *E. formosa* |
|---|---|---|---|---|---|---|---|---|---|---|---|---|
| **0** | 1 | 1 | 1 | 1 | 1 | 0 | 0 | 0 | 0 | 0 | 0 | 0 |
| **1** | 1 | 1 | 1 | 1 | 1 | 0 | 0 | 1 | 0 | 0 | 0 | 0 |
| **2** | 1 | 1 | 1 | 1 | 1 | 0 | 0 | 0 | 1 | 0 | 0 | 0 |
| **3** | 1 | 1 | 1 | 1 | 1 | 0 | 0 | 0 | 0 | 0 | 1 | 0 |
| **4** | 1 | 1 | 1 | 1 | 1 | 0 | 0 | 0 | 1 | 0 | 1 | 1 |
| **5** | 1 | 1 | 1 | 1 | 1 | 0 | 0 | 1 | 0 | 1 | 1 | 0 |
| **6** | 1 | 1 | 1 | 1 | 1 | 1 | 0 | 0 | 3 | 2 | 0 | 0 |
| **7** | 1 | 1 | 1 | 1 | 1 | 0 | 0 | 0 | 2 | 2 | 0 | 9 |
| **8** | 1 | 1 | 1 | 1 | 1 | 0 | 1 | 0 | 1 | 3 | 1 | 0 |
| **9** | 1 | 1 | 1 | 1 | 1 | 1 | 0 | 1 | 5 | 0 | 3 | 0 |
| **10** | 1 | 1 | 1 | 1 | 1 | 1 | 0 | 0 | 1 | 5 | 1 | 0 |
| **11** | 1 | 1 | 1 | 1 | 1 | 0 | 1 | 2 | 0 | 5 | 1 | 0 |
| **12** | 1 | 1 | 1 | 1 | 1 | 0 | 1 | 5 | 5 | 3 | 2 | 0 |
| **13** | 1 | 1 | 1 | 1 | 1 | 0 | 0 | 3 | 5 | 3 | 2 | 2 |
| **14** | 1 | 1 | 1 | 1 | 1 | 2 | 0 | 1 | 3 | 5 | 1 | 0 |
| **15** | 1 | 1 | 1 | 1 | 1 | 1 | 1 | 2 | 5 | 2 | 1 | 0 |
| **16** | 1 | 1 | 1 | 1 | 1 | 0 | 1 | 3 | 8 | 2 | 3 | 1 |

***Table S5. Summary of COI sequencing.*** *Summary statistics of the sequencing performed for the three experiments.*

| SUMMARY | Total COI sequences; Filtered. merged and chimeras removed | Minimum per sample | Average per sample | Median per sample | Maximum per sample |
|---|---|---|---|---|---|
| **Experiment I** | 696253 | 1111 | 9806.380282 | 9024 | 20568 |
| **Experiment II** | 768752 | 1200 | 7391.846154 | 6931.5 | 18907 |
| **Experiment III** | 1790236 | 448 | 7665.320513 | 5564.5 | 17012 |
| **AVERAGE OVERALL** | | | 7572.826087 | | |
| **SD** | | | 3869.42211 | | |
